# Machine learning approaches to identify core and dispensable genes in pangenomes

**DOI:** 10.1101/2021.03.22.436446

**Authors:** Alan E. Yocca, Patrick P. Edger

## Abstract

A gene in a given taxonomic group is either present in every individual (core), or absent in at least a single individual (dispensable). Previous pangenomic studies have identified certain functional differences between core and dispensable genes. However, identifying if a gene belongs to the core or dispensable portion of the genome requires the construction of a pangenome, which involves sequencing the genomes of many individuals. Here we aim to leverage the previously characterized core and dispensable gene content for two grass species (*Brachypodium distachyon* and *Oryza sativa*) to construct a machine learning model capable of accurately classifying genes as core or dispensable using only a single annotated reference genome. Such a model may mitigate the need for pangenome construction, an expensive hurdle especially in orphan crops which often lack the adequate genomic resources.

## Introduction

Reference genome assemblies contain information specific only to the individual of the species sequenced to create the assembly. They lack genomic regions present in other individuals of that species. Recently, the widespread adoption of pangenomics enabled characterization of the gene content diversity present in a species (Golicz et al. 2016; Montenegro et al. 2017; Gordon et al. 2017; Hurgobin et al. 2018; Ou et al. 2018; Li et al. 2014; Zhou et al. 2017; Gao et al. 2019; Hübner et al. 2019; Yu et al. 2019; Lin et al. 2014; W. Wang et al. 2018). The term pangenome was first coined in 2005, referring to collections of sequences across different strains of microorganisms (Tettelin et al. 2005). This early work built upon the observation that genes often display presence-absence variation (PAV) across different strains. Genes present in every individual of a taxonomic group are called core genes, while genes absent in at least a single individual are called dispensable genes.

The generation of a pangenome allows us to determine if a particular gene in each reference assembly is either core or dispensable according to their presence or absence in individuals used to construct the pangenome. Additionally, we know there are both qualitative and quantitative differences between core and dispensable genes. For example, in plants, core genes are often associated with essential metabolic processes while dispensable genes are associated with adaptive functions (e.g. stress responses; Danilevicz et al. 2020). Previous work also demonstrated dispensable genes exhibit higher rates of polymorphism than core genes (Gordon et al. 2017; Li et al. 2014; Hurgobin et al. 2018; W. Wang et al. 2018). This framework is analogous to a binary classification problem, one potentially addressed by machine learning.

We use the term “machine learning” to refer to the application of computer algorithms to classify observations based on previous information. Machine learning has increasingly more often been applied to genomics research (Golicz et al. 2020). For example,machine learning has been used to predict gene expression levels from genomic sequence data (Azodi, Lloyd, and Shiu, n.d.). Machine learning has also been used in the biomedical field to diagnose disease (Kourou et al. 2015). A broad application of machine learning is executed in Deep Variant, a software tool that identifies variants based on short-read sequence alignments (Poplin et al. 2018).

Here we aim to apply machine learning algorithms to classify genes as core or dispensable in a new genome given nothing except a few simply determined characteristics of a gene. We first identify quantitative differences between core and dispensable genes in two different grass species, *Oryza sativa* and *Brachypodium distachyon,* for which high-quality pangenomes were developed (Gordon et al. 2017; W. Wang et al. 2018). Furthermore, a shared ancient polyploidization event and phylogenetic placement near many additional agronomically important species make these species befitting for our study. Then, we trained different machine learning models to differentiate between core and dispensable genes based on yet to be determined differences. Finally, we tested the feasibility of applying these trained models to species not used to train our models.

## Results

### Differences between core and dispensable genes

Previous pangenome studies have revealed that there are some functional differences between core and dispensable genes (Danilevicz et al. 2020). We investigated quantitative differences between gene models in reference genomes listed as core or dispensable according to two previous pangenomes. Wang et al. analyzed gene PAV across 453 *Oryza sativa* accessions (W. Wang et al. 2018). They found that roughly 58% of genes in the reference assembly were present in each of the 453 accessions. Gordon et al. generated full *de novo* assemblies for 54 *Brachypodium distachyon* accessions (Gordon et al. 2017). They discovered that roughly 30% of genes in the reference genome were present in each accession. These two systems provide us with independent pangenome assemblies to train and test prediction models.

We observe quantitative differences for various features between the gene models of core and dispensable genes (Fig 1). Interestingly, we observe a bias for higher guanine-cytosine base-pair (GC) percentage in dispensable genes. As observed before, gene models in grass genomes have a bimodal GC percentage distribution (Clément et al. 2014; McKain et al. 2016). Other quantitative gene feature differences between core and dispensable gene models include: gene length, exon count, intron count, exon length, and intron length (Figure 1; Fig S1).

**Figure 1:**
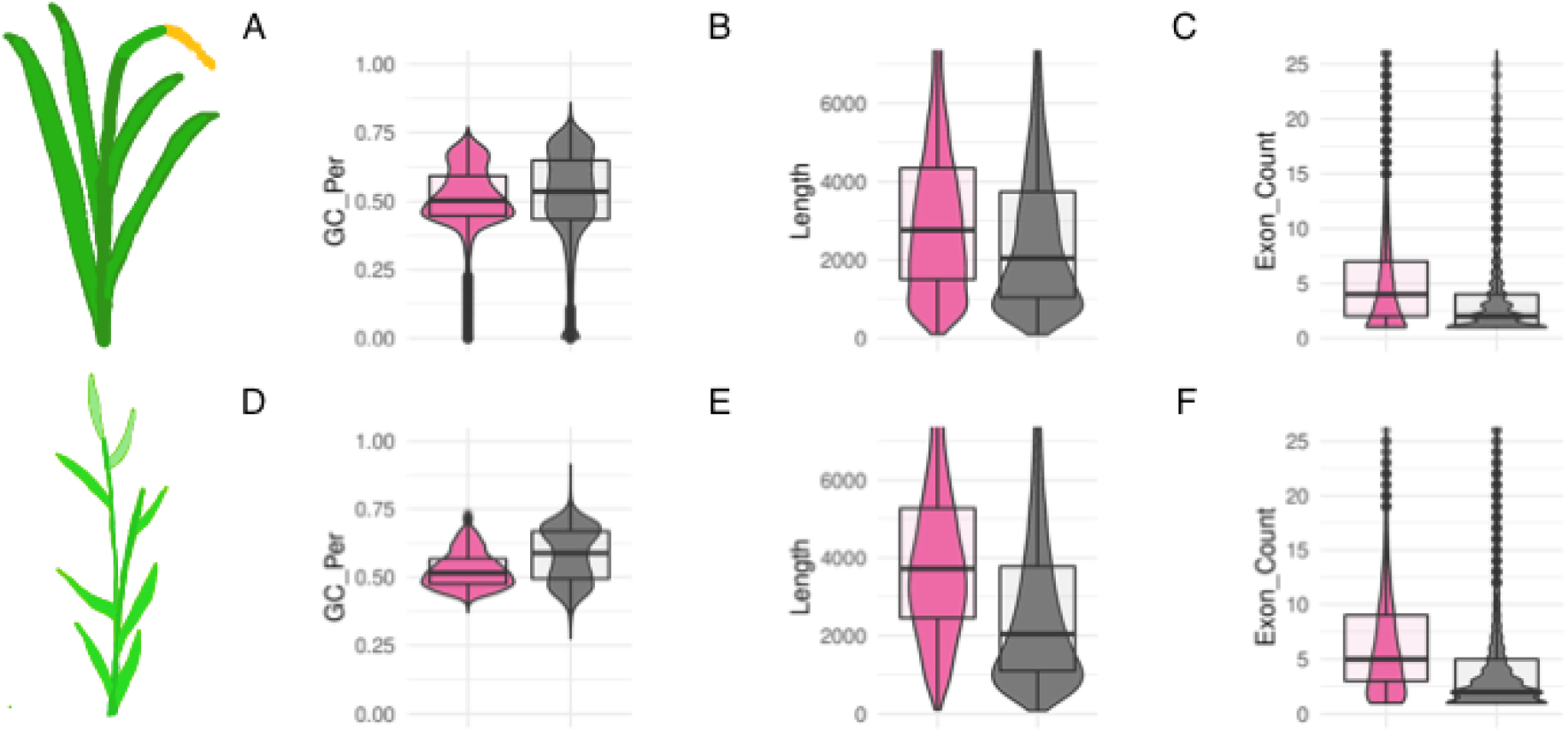
Quantitative differences in core (pink) and dispensable (grey) gene models for both *O. sativa* (A,B,C) and *B. distachyon* (D, E, F). Features are as follows: (A,D) Guanine-Cytosine percentage, (B,E) gene length measured from annotated transcription start site to transcription end site, and (C,F) number of exons.

As dispensable genes are absent in at least one individual in a taxonomic group, we hypothesize core and dispensable genes may evolve differently. One signature of past selection is the ratio of non-synonymous to synonymous substitutions (Ka/Ks ratio). Previous pangenome studies have reported a variety of approaches comparing non-synonymous and synonymous substitution rates. Consistently, they find higher non-synonymous substitution rates, as well as elevated Ka/Ks ratios for dispensable genes compared to core genes implying greater positive selection acts on dispensable genes (W. Wang et al. 2018; Gordon et al. 2017; Golicz et al. 2016; Li et al. 2014; Pinosio et al. 2016; Hurgobin et al. 2018).

There are two different ways to calculate Ka/Ks values: (1) alignments between intragenomic paralogs or (2) alignments between orthologs across species. Both methods were applied here. We found 88.25% of *O. sativa* genes have a paralog, while 52.4% of genes have an ortholog with a *B. distachyon* gene. For *B. distachyon*, these values were 87.1% and 67.4% for paralogs and orthologs to a *O. sativa* gene respectively. As reported in previous studies, Ka/Ks ratio distributions were higher for dispensable genes than for core genes (Fig S1).

A consistent observation in pangenomic studies is that dispensable genes are enriched with functions associated with biotic and abiotic stress response. Therefore, we wanted to incorporate these underlying sequence differences in our models. We considered some sort of quantitative measure of gene-ontology (GO) term similarity. However, given that the primary goal of our study is to develop a machine learning approach that may be suitable for orphan crops and lineages for which functional annotations are likely absent, we excluded GO terms for training our models. Please see other studies that have incorporated GO term differences into machine learning models (Cusack et al., n.d.).

In an attempt to account for sequence differences between core and dispensable genes without the onus of missing data, we investigated the proportion of all possible dinucleotides as a feature. This measure has a value for each gene, is readily available, and may allow our machine learning models to learn differences in underlying sequence between core and dispensable gene functions. Indeed, we observe differences in dinucleotide proportions between core and dispensable genes, suggesting this information may contribute towards core and dispensable gene differentiation (Figure S2).

### Are there differences between core and dispensable genes in relation to duplication type?

Gene duplications have played a major role in shaping the gene content in eukaryotic genomes and have contributed to the evolution of novel traits (Ohno 1970). There are multiple mechanisms of gene duplication that may exhibit differences between core and dispensable genes as reported previously in sesame (Yu et al. 2019). We used MCScanX, a toolkit for evolutionary analyses, to classify each gene in both *O. sativa* and *B. distachyon* (Figure 2) into different gene duplication classes with the intention to test whether core or dispensable genes are enriched for tandem or whole genome duplicates. McScanX provides a function called duplicate_gene_classifier that assigns genes to one of five classes: Dispersed, Proximal, Singleton, Tandem, or Whole Genome Duplicate. Dispersed duplicates are those existing > 20 genes apart from each other and not belonging to any other listed category. Proximal duplicates are paralogs located within 20 genes of each other. Singleton genes do not have a paralog. Tandem duplicates are labeled as paralogous pairs existing next to each other without any intervening genes. Whole genome duplicates are those that were derived from an ancient polyploid event (Y. Wang et al. 2012). The genomes of *B. distachyon* and *O. sativa* share the remnants of the same three ancient polyploidization events; rho, sigma and tau WGDs (McKain et al. 2016).

**Figure 2:**
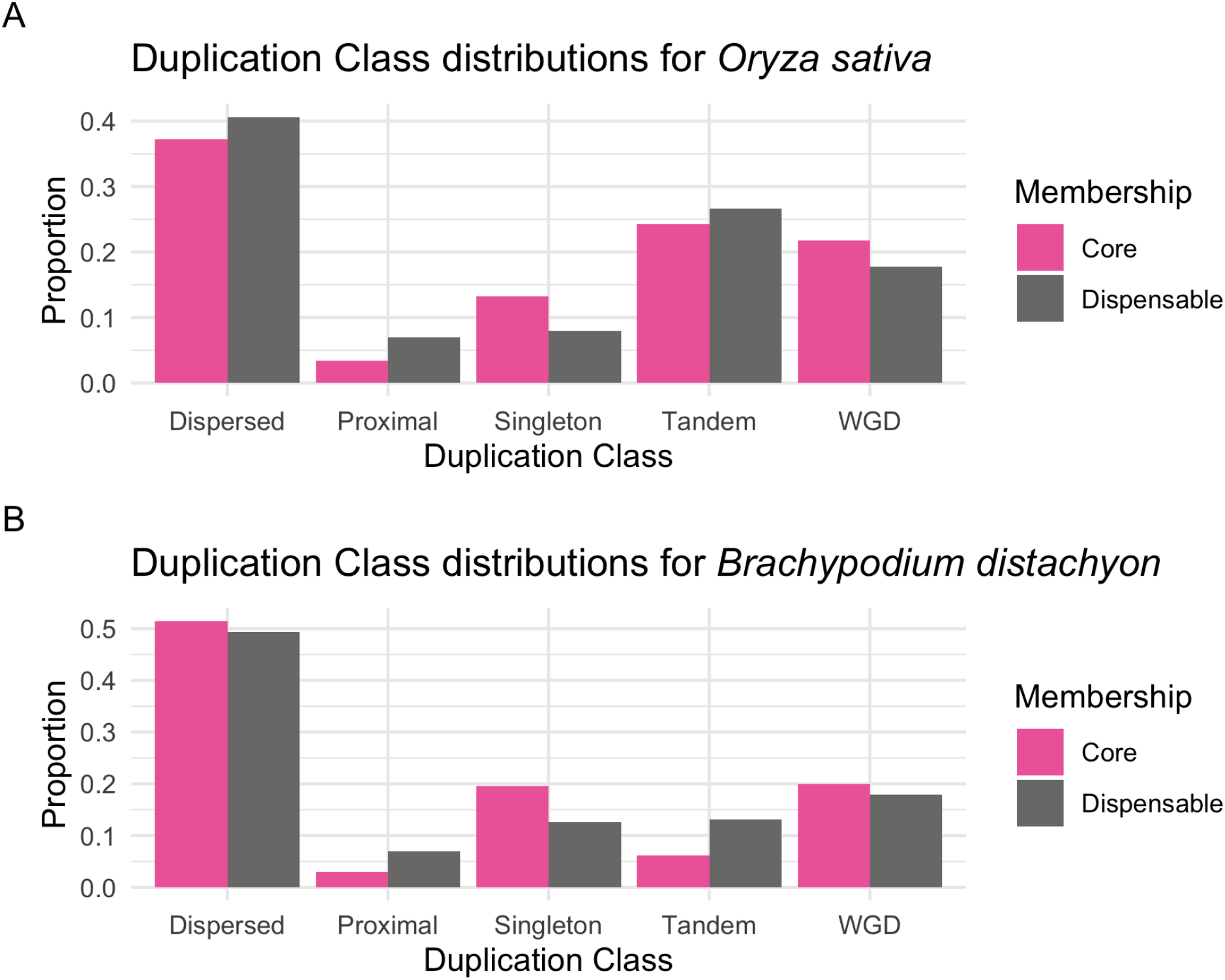
Proportion of retained duplicates by duplication class s for core and dispensable genes. Panel A depicts differences for gene models in the *O. sativa* reference genome. Panel B depicts differences for gene models in the *B. distachyon* reference genome.

We find nearly the same pattern for both *B. distachyon* and *O. sativa*. All differences between core and dispensable genes are significant (z-test p-value < 0.01). Dispensable genes contain a larger proportion of proximal and tandem duplicates, while core genes contain a larger proportion of whole-genome duplicates and single-copy genes. This pattern, combined with functional enrichment differences, is consistent with the Gene Balance Hypothesis (M. Freeling 2008). *O. sativa* and *B. distachyon* differ in which class contains a larger proportion of dispersed duplicates.

### Machine Learning methods

We tested three separate Machine Learning (ML) methods: Support-Vector Machine, Gaussian Naive-Bayes, and Random Forest. These three approaches encompass different classification techniques. Utilization of all three techniques allow us to robustly investigate our potential to differentiate between core and dispensable genes.

Detailed descriptions of these methods can be found elsewhere (Vapnik 1995; Hand and Yu 2001; Surhone, Tennoe, and Henssonow 2010). The Support-Vector Machine (SVM) classifier shapes our data in multi-dimensional space. It then searches for a vector through that space which best separates our two classes, in our case core and dispensable genes. Therefore, when given a new gene to classify as core or dispensable, it plots that gene’s values in the same multidimensional space and classifies the new gene according to the created vector of best separation. Gaussian Naive-Bayes (GNB) takes each feature independently and assumes the feature values follow a gaussian distribution whose midpoint represents the difference between the two classes. With this distribution for each feature, when given a new value, it can assign a probability of either class to that value. Summing these probabilities across all features, this classifier determines the new gene’s class by the probability of belonging to either class given its feature values. The Random Forest (RF) classifier creates a forest of decision trees. A decision tree is similar to a flowchart where different paths are taken at each node. Nodes in these trees represent values of a feature that will send a new gene along different classification paths depending on their values. Random Forest is a common method used in classification and is capable of learning high order and non-linear associations in the classification data.

### Model training and assessment

We could train our models using all of our data, however testing on that same data may allow the models to effectively “memorize” certain genes’ feature values, a phenomenon called “over-training”. These models will perform poorly on unseen data. To prevent over-training, we only want to train our models on a subset of our data. That subset may not reflect all the patterns in our data. To ensure our model performance is not affected by a non-representative subset of our data, we apply a k-fold cross validation approach. We split our data into k equally sized subsets (k = 10 in our case). For each subset, we train our models on all other subsets, and test it on the remaining subset. Training and testing separately on all 10 subsets allows for a robust assessment of the model’s performance.

There are several different metrics to test model performance. We focus on accuracy and AUC-ROC. Accuracy is simply the proportion of correctly classified genes in the testing subset. AUC-ROC stands for Area Under the Curve of the Receiver Operator Characteristic. This measure incorporates true positive classification rate and false positive classification rate. If the model is a random guesser, it will achieve an AUC-ROC score of 0.5 for binary classification problems. A perfect AUC-ROC score is 1 where genes are always correctly classified. This value allows us to better assess our true and false positive rates.

### La reveal magnifico

We trained and tested all three models for both species. The results are presented in Table 1. Values correspond to averages across all k-folds. Overall, all models performed better than random expectations. This indicates we are able to learn differences between core and dispensable genes in different species and classify unseen genes as core or dispensable. Model performance is overall better in *B. distachyon* than in *O. sativa*. The Random Forest model out performed other models in terms of both accuracy and AUC-ROC.

**Table 1:** Preliminary model assessment. Values are means and ranges are the standard deviation from 10 cross-fold validations. Abbreviations are as follows: Support Vector Machine (SVM), Gaussian Naive Bayes (GNB), Random Forest (RF), and Area Under the Curve for the Receiver Operator Curve (AUC-ROC).

| Species | Model | Accuracy | AUC-ROC |
| --- | --- | --- | --- |
| *Oryza sativa* | SVM | 0.684 +/- 0.018 | 0.738 +/- 0.025 |
| *Oryza sativa* | GNB | 0.651 +/- 0.019 | 0.704 +/- 0.025 |
| *Oryza sativa* | RF | 0.700 +/- 0.017 | 0.763 +/- 0.023 |
| *Brachypodium distachyon* | SVM | 0.762 +/- 0.011 | 0.788 +/- 0.015 |
| *Brachypodium distachyon* | GNB | 0.704 +/- 0.013 | 0.784 +/- 0.012 |
| *Brachypodium distachyon* | RF | 0.800 +/- 0.010 | 0.857 +/- 0.011 |

### What features are most important for classifying core and dispensable genes?

Not every feature is likely to contribute equally to model performance. To determine which features are most important for differentiating between core and dispensable genes, we performed recursive feature elimination, one of several strategies of feature selection. In recursive feature elimination, we train our model using all features, and the least important feature is eliminated. After elimination, we re-train our model and perform the same operation. We measure model accuracy at each step. By viewing the accuracy as features are eliminated, we can select the combination of features that provides us with the greatest accuracies. These curves for both the Support Vector Classifier and Random Forest models are shown in Fig S4. We find using all features results in the highest model performance compared to excluding low performing features.

The Random Forest and Support Vector Classifier models explicitly provide relative feature importance scores. These scores reflect how much relative weight each feature contributes to the final prediction. Relative feature importance scores are shown in Fig S5. Overall, several measures related to GC percentage stand out as large contributors to final predictions. Comparing the GC percentage between core and dispensable genes in both *B. distachyon* and *O. sativa* reveal striking differences in this measure (Fig 1A,1D). Therefore, we believe GC percentage is an important distinguishing character between core and dispensable genes in these grass species.

### How does a model trained on one species perform on the other?

We trained our models on one species and tested it on the other species. In this instance, it is important to consider the proportion of core and dispensable genes in each reference genome. A model trained on a species with 70% core genes will anticipate 70% of the testing data to be core as well. Additionally, if 70% of genes in a reference genome are core, a model can obtain an accuracy of 70% simply by predicting every case to be core. This emphasizes the importance of measures other than accuracy alone to evaluate models such as AUC-ROC.

The proportion of core genes in a reference genome is variable across lineages (Golicz et al. 2020). To account for these differences, we both (1) test balanced training and testing data, as well as (2) measure AUC-ROC rather than accuracy. To balance our datasets, we subset the majority class to the size of the minority class. For example, in *B. distachyon*, we find ~30% (n = 9,308) of genes are core out of all used to train our models (n = 31,679). We balance this data by subsetting 9,308 core genes and train our models on only 18,616 genes rather than the 31,679 total gene models.

As expected, model performance is worse cross-species than within species (Table S1). Importantly, Random Forest model performance training with *B. distachyon* gene models then testing on *O. sativa* gene models performed worse than random expectation (AUC-ROC = 0.4853). However, all other combinations outperformed random expectation.

There are two possible reasons for decreased model performance. First, there may be lineage specific differences between core and dispensable genes. Indeed, we visualize these differences by plotting gene model distributions on the same axes comparing *O. sativa* and *B. distachyon* (Fig S1). Second, though we attempted to correct for it, our models could be overtrained on our data.

We especially notice differences between both species in relation to their Ka and Ks distributions. To test if these differences were responsible for poor model performance, we trained and tested our models without any Ka or Ks values. Interestingly, Random Forest model performance was dramatically improved when excluding these features (Table 2).

**Table 2:** Table 2 shows cross-species model performance (AUC-ROC) compared to intra-specific model performance. Features related to Ka and Ks were excluded from models to produce these results.

| Training data | Testing data | Model | AUC-ROC | AUC-ROC self |
| --- | --- | --- | --- | --- |
| *Oryza sativa* | *Brachypodium distachyon* | SVC | 0.6953 | 0.7354 +/- 0.026 |
| *Oryza sativa* | *Brachypodium distachyon* | GNB | 0.6892 | 0.6924 +/- 0.027 |
| *Oryza sativa* | *Brachypodium distachyon* | RFT | 0.6646 | 0.7313 +/- 0.022 |
| *Brachypodium distachyon* | *Oryza sativa* | SVC | 0.6255 | 0.7481 +/- 0.015 |
| *Brachypodium distachyon* | *Oryza sativa* | GNB | 0.6519 | 0.7485 +/- 0.012 |
| *Brachypodium distachyon* | *Oryza sativa* | RFT | 0.5978 | 0.8059 +/- 0.012 |

## Discussion

In this study, we tested the efficacy of training machine learning models to differentiate between core and dispensable genes in a single genome based on various gene features. We first sought to characterize differences between core and dispensable genes in two species (*Brachypodium distachyon* and *Oryza sativa*) with available pangenomes. Determining the origin of dispensable genes is beyond the scope of this study. However, our observations of shorter dispensable genes with fewer exons suggest a fraction of them may have arisen *de novo*. This observation is consistent with the hypothesis that short sequences are more likely to gain genic functions from previously non-coding DNA than longer sequences, and reports of *de novo* gene origin in yeast and *Drosophila* (Carvunis et al. 2012; Siepel 2009; Fig 1). However, gene duplications are likely still the predominant source of new genes.

We observed differences in the evolutionary origin and evolutionary signatures between core and dispensable genes. In addition to differences in length and intron count, we investigated non-synonymous to synonymous substitution rate (Ka/Ks) as well as duplication type differences between core and dispensable genes. We calculated non-synonymous and synonymous substitution rates in PAML (Yang 2007) using two different approaches. First, we aligned orthologs between *B. distachyon* and *O. sativa*. Second, we aligned paralogs within each of the two species. The differences in distributions between core and dispensable genes for paralogs and orthologs yielded similar results for both species. As shown previously, elevated rates of Ka/Ks observed in dispensable genes relative to core genes imply higher rates of positive selection on these genes and possibly the evolution of novel gene functions (matching the hypothesis outlined by Susumu Ohno; Ohno 1970).

The duplication history of each gene in both genomes was examined. Previous studies suggested that core and dispensable genes are enriched with different classes of gene duplicates (Yu et al. 2019). The observed gene duplication differences are consistent with hypotheses of dosage sensitive genes, as outlined by the Gene Balance Hypothesis (James A. Birchler and Veitia 2007, 2012). If core genes encode for more essential cellular functions, which are known to be enriched with highly dosage sensitive genes (Michael Freeling 2009), they would contain a higher proportion of retained duplicates from ancient whole genome duplications. Dosage sensitive genes must retain duplicates from ancient polyploid events to maintain proper stoichiometry in macromolecular complexes and gene networks (J. A. Birchler et al. 2001). This skewed pattern for retained whole genome duplicates was observed for core genes in both the rice and *Brachypodium* genomes. Similarly, previous studies have suggested that certain single copy genes encode for essential functions, including organellar-nuclear interactions (Edger and Pires 2009), and must remain in single copy due to gene dosage constraints (De Smet et al. 2013; Tasdighian et al. 2017). Our analyses also show that core genes are enriched with a greater number of single copy genes.

Dispensable genes, on the other hand, are enriched with more adaptive functions. Adaptive genes tend to be more poorly connected and, thus, are heavily skewed towards being more dosage insensitive (Rizzon, Ponger, and Gaut 2006). Dosage insensitive genes are known to be enriched with tandem duplicated genes (M. Freeling 2008; James A. Birchler and Veitia 2012). Similarly, the dispensable gene content of the pangenome, as shown in this study, is enriched with tandem duplicates. Thus, tandem duplication appears to be the prominent mechanism giving rise to new dispensable genes. In summary, core genes contain a higher proportion of retained duplicates from whole genome duplications and single copy genes, while dispensable genes contain a higher proportion of retained tandem duplicates.

Applying three separate machine learning models revealed similar results. We are able to differentiate between core and dispensable genes better than random, yet with imperfect accuracy. Our three models displayed different performances likely due to differences in the distributions of the data. For example, we believe the Gaussian Naive Bayes model performed the worst since it fits each feature to a normal distribution when the distribution of each feature is not normally distributed. Additionally, it gives equal weight to each feature in the decision making process. Our other two models demonstrate each feature contributes a unique amount to the final prediction resulting in worse performance for the Gaussian Naive Bayes model. The Random Forest model out performed the Support Vector Machine model. This observation is consistent with other applications of machine learning which demonstrate Random Forest often outperforms other models (except Cusack et al., n.d.). We recommend using multiple models on applications of classifying genes as core or dispensable in the future.

Model performance performed better within than across species. Previous applications of machine learning across species yielded similar results (Lee, Karchin, and Beer 2011; Chen, Fish, and Capra 2018; Kelley, n.d.; Mejía-Guerra and Buckler 2019). For example, Meng et al. trained machine learning models to identify cold-responsive genes across a few different grass species (Meng et al. 2021). Consistent with our results, models performed best when trained and tested in the same species. Notably, they mentioned shared phenotypes are better indicators of cross-species model performance than ancestry. Therefore, perhaps our cross species model performances would be improved by testing in not only phylogenetically closer taxa, but also those exhibiting similar phenotypic and perhaps pangenome characteristics.

A potential application of these models is to classify genes as core or dispensable in a new species without the costly construction of a pangenome. While our models perform better than random guessing, our accuracy rates are insufficient to substitute for pangenome construction for many downstream applications. However, if ~70% accuracy is all that is required, perhaps in the case of developing a genotyping array that consists of largely core genes for guiding breeding efforts, this strategy may likely suffice. We recommend training a model on a species as closely related to the study species as possible. Therefore, we advocate for a community-wide effort for pangenome construction of strategically phylogenetically placed taxa. Broad pangenome development will further increase our understanding of not only what combination of features differentiate core and dispensable genes, but also on various topics ranging from better understanding the evolutionary dynamics of gene families to genotype-phenotype associations.

## Methods

### Core and dispensable gene annotations

#### Oryza sativa

We obtained a matrix of gene presence and absence from https://figshare.com/articles/dataset/Gene_presence_absence_variations_of_453_rice_accessions/5103769 (last accessed 03-08-2021) taken from a recent publication (Wang et al. 2018). This study analyzed short read sequencing from <3,000 rice accessions, yet included gene PAV for 453 accessions (the authors selected these accessions for “sequencing depths > 20× and mapping depths > 15×”).

The gene PAV matrix file is GenePAV.matrix.txt. It provides PAV information coded in binary (1 for presence, 0 for absence). Locus identifiers are provided according to the annotation downloaded below (eg Os01g0100100). Therefore, specific transcript information is unavailable. This potentially affects a few gene measures such as exon/intron count and gene length. We take information for the longest listed transcript for each locus. We only consider loci with an available annotation in the IRGSP-1.0 rice annotation release. Therefore, we have PAV information for 35,633 genes with a locus identifier. Using this matrix, we defined core genes as those present in each of the 453 accessions in the matrix. Dispensable genes are those absent in at least a single accession.

#### Brachypodium distachyon

*B. distachyon* core and dispensable gene information was downloaded from https://genome.jgi.doe.gov/portal/pages/dynamicOrganismDownload.jsf?organism=BrachyPan (last accessed 03-08-2021; JGI login required).

We found information from 54 brachypodium lines in accordance with Supplementary Table S1 from Gordon et al. (Gordon et al. 2017). We then created a PAV matrix for every locus in the reference genotype Bd21, again selecting the longest transcript. Again, we define core genes as those present in all individuals and dispensable genes as those missing in at least a single accession.

#### Genome and annotation versions

We used the same genome and gene annotation versions as used in the pangenome studies from which we gathered core and dispensable annotations. For *O. sativa*, we collected the IGRSPv1.0 annotation here: https://rapdb.dna.affrc.go.jp/download/irgsp1.html (last accessed 03-08-2021). For *B. distachyon*, we collected the Brachypodiumv2.1 annotation from JGI.

### Feature calculations

#### Quantitative gene features

All gene features were gathered using the scripts ‘annotate_core_genes_osat_nested.py’ and ‘annotate_core_genes_bdis.py’ for *O. sativa* and *B. distachyon* respectively. We calculated the following features: gene length (TSS to TES), exon count, intron count, intron length, exon length, guanine-cytosine percentage, and the proportion of all possible dinucleotide pairs (AA, AT, AG, AC, TA, TT, TG, TC, GA, GT, GG, GC, CA, CT, CG, and CC).

#### Duplication type

The duplication type for each gene was determined using the MCScanX duplicate_gene_classifier function. Genes were assigned to one of five classes: Dispersed, Proximal, Singleton, Tandem, or Whole Genome Duplicate. Dispersed duplicates are those existing > 20 genes apart from each other and not belonging to any other listed category.

Proximal duplicates are paralogs located within 20 genes of each other. Singleton genes do not have a paralog. Tandem duplicates are labeled as paralogous pairs existing next to each other without any intervening genes. Whole genome duplicates are those labeled as anchor genes, in other words those which scaffold intragenomic collinear blocks (Y. Wang et al. 2012). These anchor genes are hypothesized to have been duplicated by a polyploidization event. These classes were coded numerically as 0, 1, 2, 3, and 4 respectively.

#### Substitution rate calculations

We calculated non-synonymous and synonymous substitution rates in PAML using two different comparisons. First, we aligned orthologs between *B. distachyon* and *O. sativa*. Second, we aligned paralogs within each of the two species. PAML was run through a custom pipeline available here: https://github.com/Aeyocca/ka_ks_pipe (last accessed 03-08-2021). As not every gene model has a paralog and ortholog, there were missing values. We chose to code these missing values as an arbitrary number (99) the model could recognize as different from a potential true value.

### Model training and evaluation

Machine learning models were trained and evaluated using the scikit learn toolkit in the python programming language (Pedregosa et al. 2011). A few scripts were used to implement these functions:

Calculation of AUC-ROC for within and cross species predictions:
  ‘osat_bdis_kfold_model_test_auc_21_02.py’ and
Creation of AUC-ROC curves for within-species 10-fold cross validation:
  ‘osat_auc_roc_curves.py’ and ‘bdis_auc_roc_curves_array.py’
Calculation of accuracy for within and cross species predictions:
  ‘osat_bdis_kfold_model_test_21_02.py’
Calculation of feature importance values for the Random Forest and Support Vector Machine models:
  ‘osat_bdis_feat_imp.py’

## Availability of supporting data

Scripts used in this study are publically available on github https://github.com/Aeyocca/Core_Prediction (last accessed 03-08-2021)

## Acknowledgements

This work was supported by Michigan State University AgBioResearch, National Science Foundation (DEB #1737898) and United States Department of Agriculture (AFRI #2019-51181-30015)). We thank members of the Edger Lab for helpful feedback.

## Supplementary Figures

**Figure S1:**
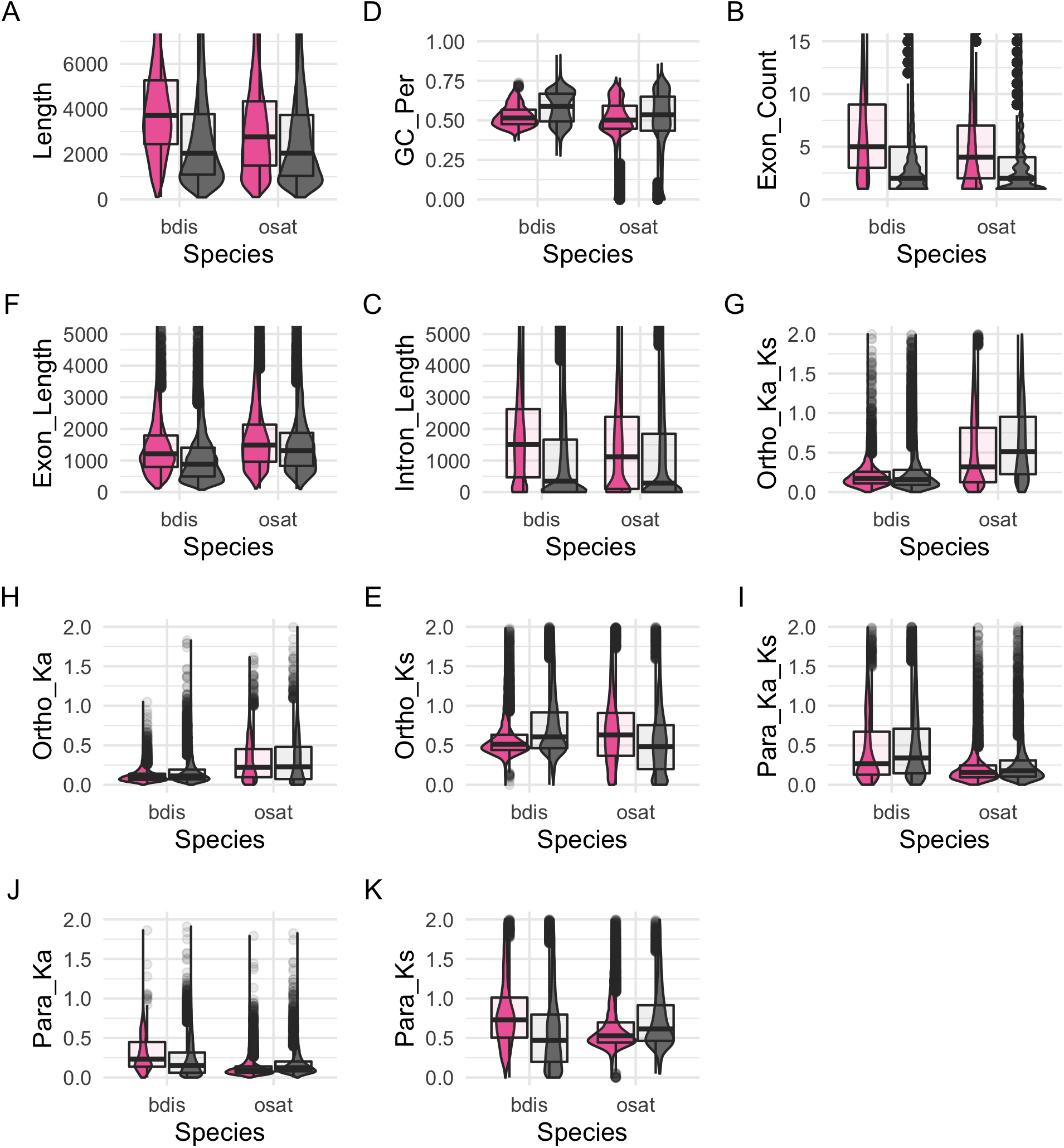
Feature distributions scaled. Here we show feature distributions of core (pink) and dispensable (grey) genes for both *Oryza sativa* (osat) and *Brachypodium distachyon* (bdis). We show these distributions on the same axes to suggest reduced cross-species model performance may be attributed to lineage specific distributions.

**Figure S2:**
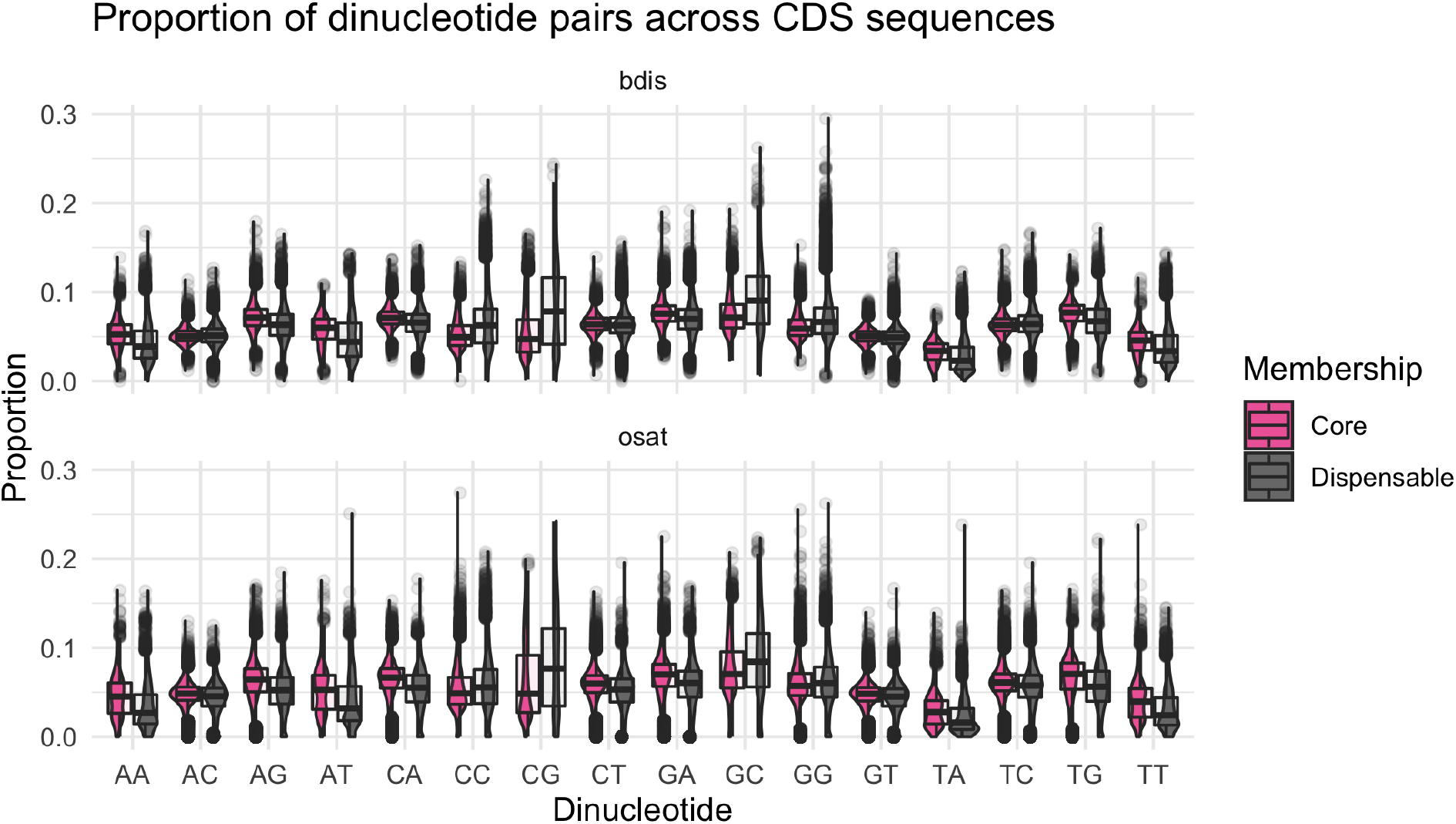
Distributions of dinucleotide proportions. We plot the proportion of dinucleotide proportions across all coding sequences (CDS) in both *Brachypodium distachyon* (bdis) and *Oryza sativa* (osat). We directly compare these distributions between core and dispensable genes.

**Figure S3:**
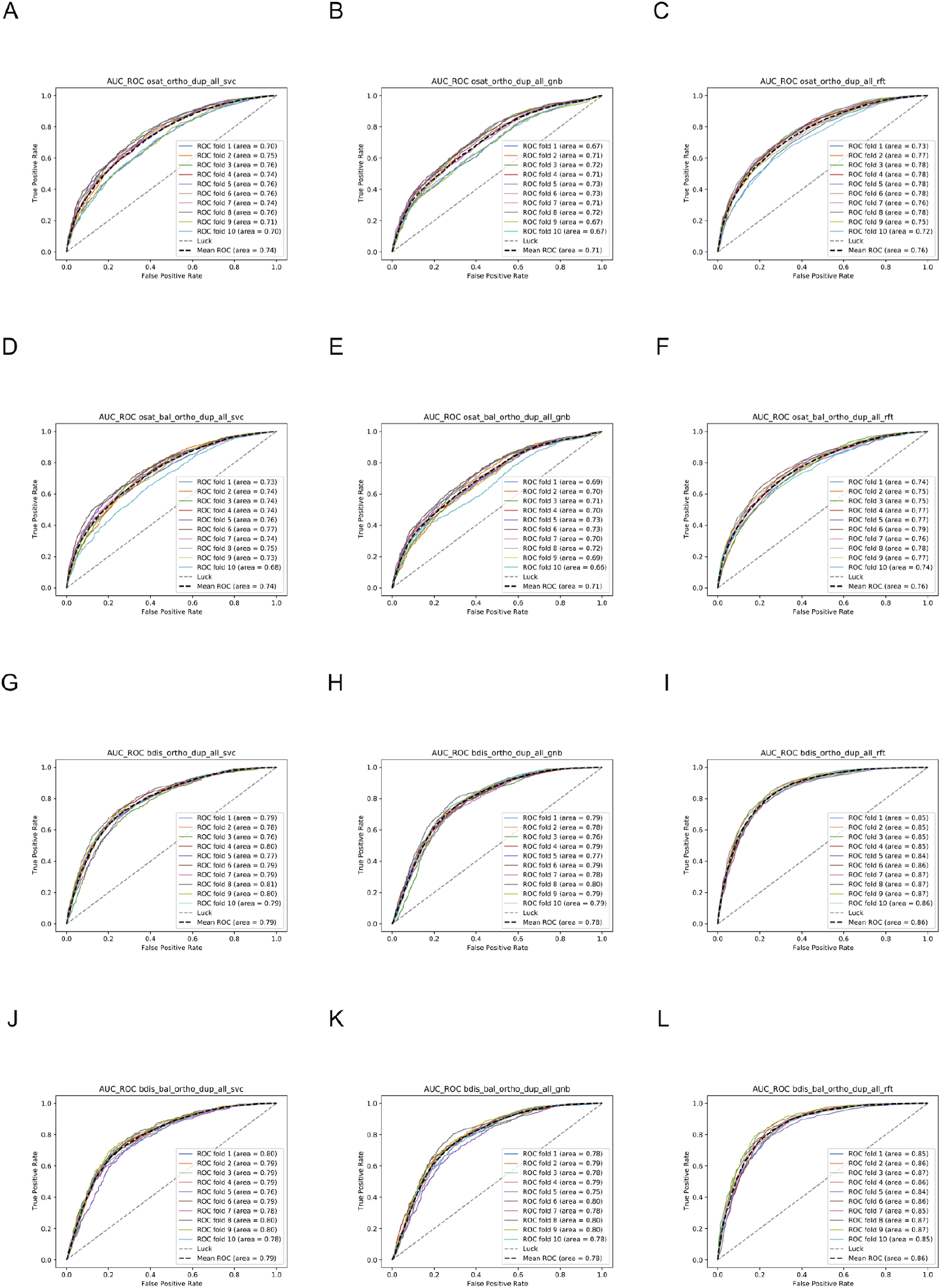
Figure S3 shows AUC-ROC graphs for different models and data subsets. Each colored line corresponds to the ROC curve for each of 10 subsets. The dotted line shows the average across all 10 subsets. The diagonal line corresponds to the random expectation of 0.5.

**Figure S4:**
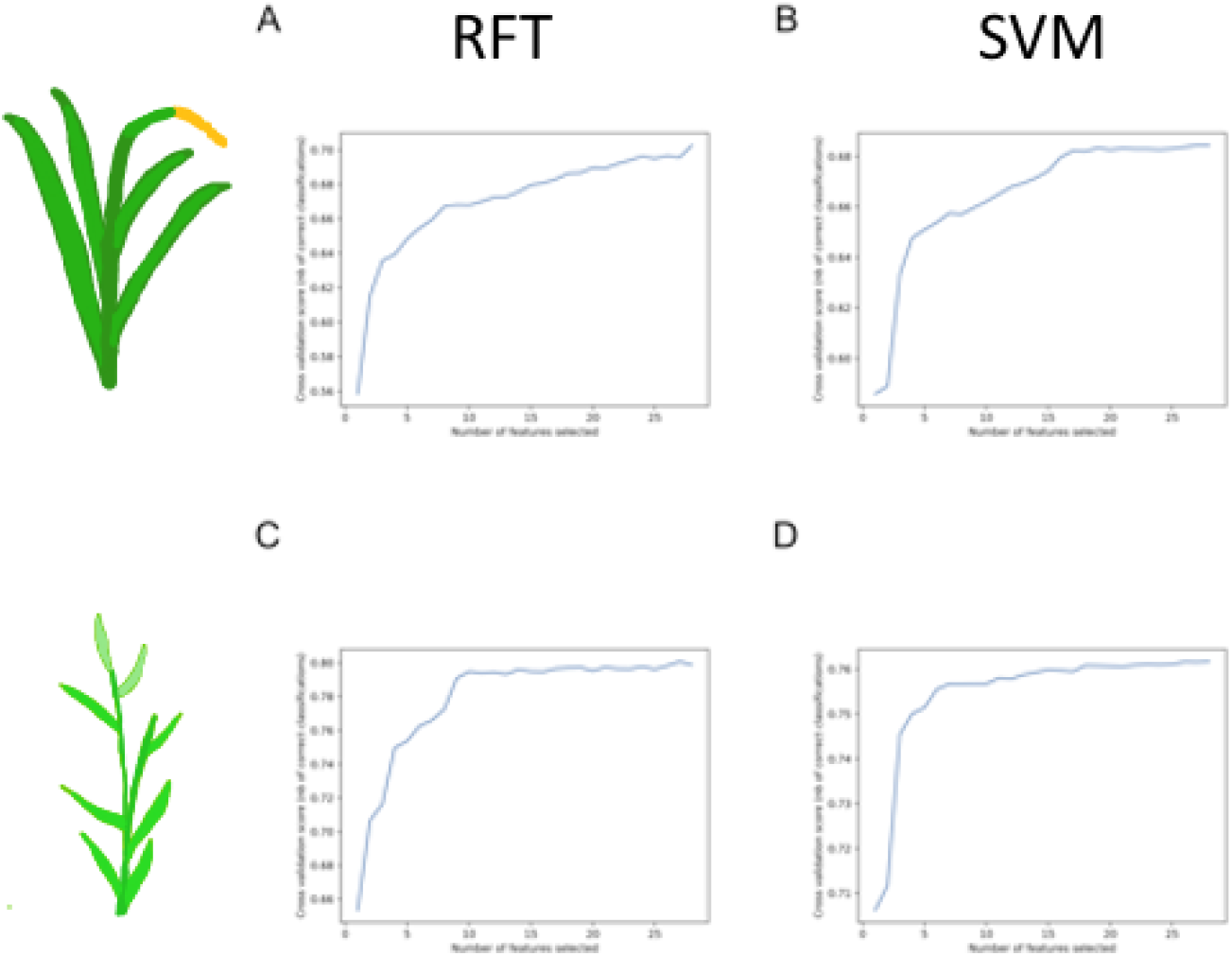
Figure S4 shows the recursive feature elimination curves for the Random Forest (RFT) and Support Vector Machine (SVM) models for *O. sativa* and *B. distachyon*.

**Figure S5:**
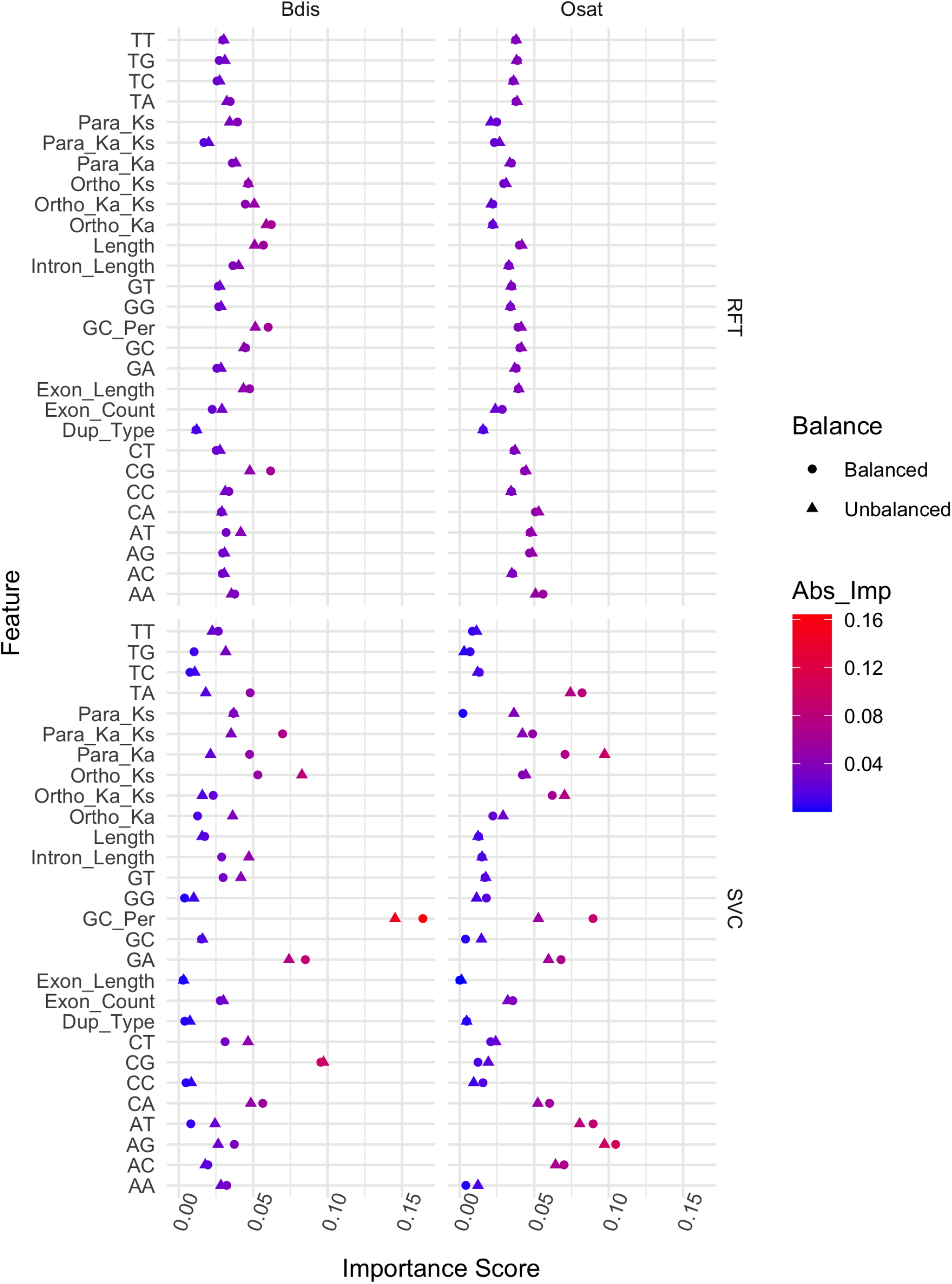
Figure S5 shows the feature importance scores for the Support Vector Machine (SVM) and Random Forest (RFT) model trained using either balanced (circles) or unbalanced (triangle) data from *Oryza sativa* (Osat) and *Brachypodium distachyon* (Bdis). The absolute values of feature importance scores are shown

